# An Intersection between Iron Availability and *Candida albicans* Invasive Filamentation

**DOI:** 10.1101/2021.09.15.460572

**Authors:** Ashlee Junier, Anne Weeks, Ysabella Alcaraz, Carol A. Kumamoto

## Abstract

*Candida albicans* filamentation, the ability to convert from oval yeast cells to elongated hyphal cells, is a key factor in its pathogenesis. Previous work has shown that the integral membrane protein Dfi1 is required for filamentation in cells grown in contact with a semi-solid surface. Investigations into the downstream targets of the Dfi1 pathway revealed potential links to two transcription factors – Sef1 and Czf1. Sef1 regulates iron uptake and iron utilization genes in low iron conditions, leading us to hypothesize that there exists a link between iron availability and contact-dependent invasive filamentation. Here, we showed that Sef1 was not required for contact dependent filamentation, but it was required for WT expression levels of a number of genes during growth in contact conditions. Czf1 is required for contact-dependent filamentation and for WT levels of expression of several genes. Constitutive expression and activation of either Sef1 or Czf1 individually in a *dfi1* null strain resulted in a complete rescue of the *dfi1* null filamentation defect. Because Sef1 is normally activated in low-iron environments, we embedded WT and *dfi1* null cells in iron-free agar medium supplemented with various concentrations of Ferrous Ammonium Sulfate (FAS). *dfi1* null cells embedded in media with a low concentration of iron (20uM FAS) showed increased filamentation in comparison to mutant cells embedded in higher concentrations of iron (50-500uM). WT cells produced filamentous colonies in all concentrations. Together, this data indicates that Dfi1, Czf1, Sef1, and environmental iron regulate *C. albicans* contact-dependent filamentation.

**Importance:** *Candida albicans* is an opportunistic pathogen responsible for a larger proportion of candidiasis and candidemia cases than any other *Candida* species (CDC). The ability of *C. albicans* cells to invade and cause disease is linked to their ability to filament and form hyphae. Despite this, there are gaps in our knowledge of the environmental cues and intracellular signaling that triggers the switch from commensal organism to filamentous pathogen. Here we identified a link between contact-dependent filamentation and iron availability. Over the course of tissue invasion, *C. albicans* cells encounter a number of different iron microenvironments, from the iron-rich gut to iron-poor tissues. Increased expression of Sef1-depndent iron uptake genes as a result of contact-dependent signaling will promote the adaptation of *C. albicans* cells to a low iron availability environment.

## Introduction

*Candida albicans* is a human commensal organism that commonly resides in the gastrointestinal tract (1, 2). While its presence is usually benign, immunocompromised individuals may experience any of a number of diseases caused by *C. albicans*, including candidiasis and candidemia (3). The ability of *C. albicans* to transition from a commensal organism to a pathogen is largely dependent on its ability to switch from a yeast to hyphal forms (2, 4–6). This yeast-to-hyphae transition, known as filamentation, occurs in response to many cues, including changes in temperature or pH, the presence of serum, certain nutrient deficiencies, and growth in contact with a semi-solid surface (7–11).

The integral membrane protein Dfi1 has been shown to been important in contact-dependent filamentation (12). Signaling through Dfi1 during growth on agar medium results in the binding of calcium-bound calmodulin to the cytoplasmic Dfi1 tail (13). This binding leads to the phosphorylation of the MAP Kinase Cek1, setting off a phosphorylation cascade that leads to the induction of filamentation. Deletion of both alleles of *DFI1* results in a filamentation defect in cells grown on or embedded in agar medium. It has been shown that a deletion of *DFI1* also results in a reduction in lethality of *C. albicans* in the intravenously inoculated mouse model of systemic candidiasis (12, 13). Cells with a defect in *DFI1* are still able to filament in response to other cues, such as presence of serum, as demonstrated previously (12). Despite this knowledge, the downstream genetic targets of the Dfi1 pathway are still yet to be identified.

In order to successfully invade tissues and cause disease, *C. albicans* must be able to thrive in many different iron microenvironments. While in the gastrointestinal tract, the amount of available iron is relatively high, whereas in the bloodstream or tissue, iron is sequestered by the host and less available to *Candida* (14). Throughout the process of filamentation and invasion of host tissue, *C. albicans* thus encounters a change in iron availability.

In order to thrive in all of these environments, *C. albicans* has developed a network of factors that allow it to adapt to different levels of available iron. Iron uptake and utilization are primarily controlled by two transcription factors – Sef1 and Sfu1. Sef1 is responsible for upregulating iron uptake and utilization genes in environments with low iron (15). In high iron environments, iron uptake pathways are repressed. Under high iron conditions, phosphorylated Sfu1 binds to the *SEF1* promoter in the nucleus, preventing transcription, and to Sef1 protein in the cytosol, tagging Sef1 for degradation (16). When starved for iron, Sef1 becomes phosphorylated, preventing Sfu1 binding. Sef1-P can then enter the nucleus where it promotes the transcription of iron uptake and utilization genes (16). Furthermore, Sef1 has been shown to be required for virulence in a murine model (15).

Iron availability has been shown to influence filamentation during liquid and plated growth of mutants lacking the important regulator of filamentation Efg1p (17). However, effects of iron availability specific to filamentation during growth in embedded conditions via the Dfi1p pathway have not been previously described.

Here we uncover a novel connection between contact-dependent filamentation and iron availability. To identify transcriptional targets of the Dfi1 pathway, we screened for genes that were upregulated during Dfi1 pathway activation using RNAseq. Numerous members of the Sef1 regulon were identified as differentially expressed in the presence or absence of Dfi1p. Further investigations revealed that Sef1 activation is able to bypass the invasive filamentation defect of the Dfi1 null mutant and promote contact-dependent filamentation. Taken together, the results demonstrate a role for Sef1 in the induction of *C. albicans* filamentation and invasion.

## Results

### Artificial Activation of Dfi1 to Identify Downstream Targets

The Dfi1 pathway is activated during growth of *C. albicans* on a semi-solid surface (12). Colonies grown on the surface of agar contain a heterogeneous population of cells that were exposed to numerous different microenvironments. For example, some cells are exposed to the air, some are in the center of the colony, and some are in contact with the agar surface.

Because of the heterogeneous nature of colonies grown on agar, a method to activate the Dfi1 pathway artificially and more uniformly in liquid culture was developed. The approach was based on the previous observation that treating liquid cultures of *C. albicans* with the calcium ionophore A23187 in the presence of calcium activates the Dfi1 pathway because the treatment favors binding of calcium-bound calmodulin to the cytoplasmic Dfi1 tail (13).

Therefore, Ca^++^/A23187 treatment was used to activate Dfi1-dependent Cek1p activation, and thus downstream gene expression, as described in (13). Briefly, log phase cells from WT and *dfi1* null strains growing in minimal media were treated with 4µM A23187 or a vehicle control, in Ca^++^ containing medium. After 30 min of treatment, cells were harvested in RNALater. RNA was extracted from the aliquot of cells stored in RNALater, as described in Materials and Methods. RNA from three independent cultures of WT and *dfi1* cells with and without Ca^++^/A23187 treatment was sent to the Tufts University Core Facility for RNAseq analysis. The Illumina TruSeq RNA Library preparation kit was used to prepare samples for Illumina sequencing. Reads were aligned to the *Candida albicans* SC5314 genome (assembly ASM18296v3) using Bowtie and differential gene expression was analyzed using CuffDiff. A total of 6,264 genes were analyzed for each treatment group.

To identify targets of the Dfi1 pathway, the following comparisons were made: gene expression in untreated WT cells versus gene expression in WT cells treated with Ca^++^/A23187, gene expression in WT cells treated with Ca^++^/A23187 versus gene expression in *dfi1* null cells treated with Ca^++^/A23187, and gene expression in untreated WT cells versus gene expression in untreated *dfi1* null cells (Figure 1A-C). For each of these comparisons, genes that were significantly differentially expressed by 2-fold or greater were identified. This analysis resulted in identification of 207, 123, and 156 genes respectively, or 383 distinct genes. For each of these 383 genes, expression could be increased, decreased, or show no change in each of the 3 comparison groups, resulting in 27 different possible patterns of gene expression. We focused on genes that showed differential expression in 2 or more of the 3 comparisons. Analysis of the 383 genes resulted in 93 genes that were differentially regulated in 2 or more of the comparison groups. These 93 genes represented 15 distinct patterns of gene expression (Figure 1D, table 1).

**Figure 1:**
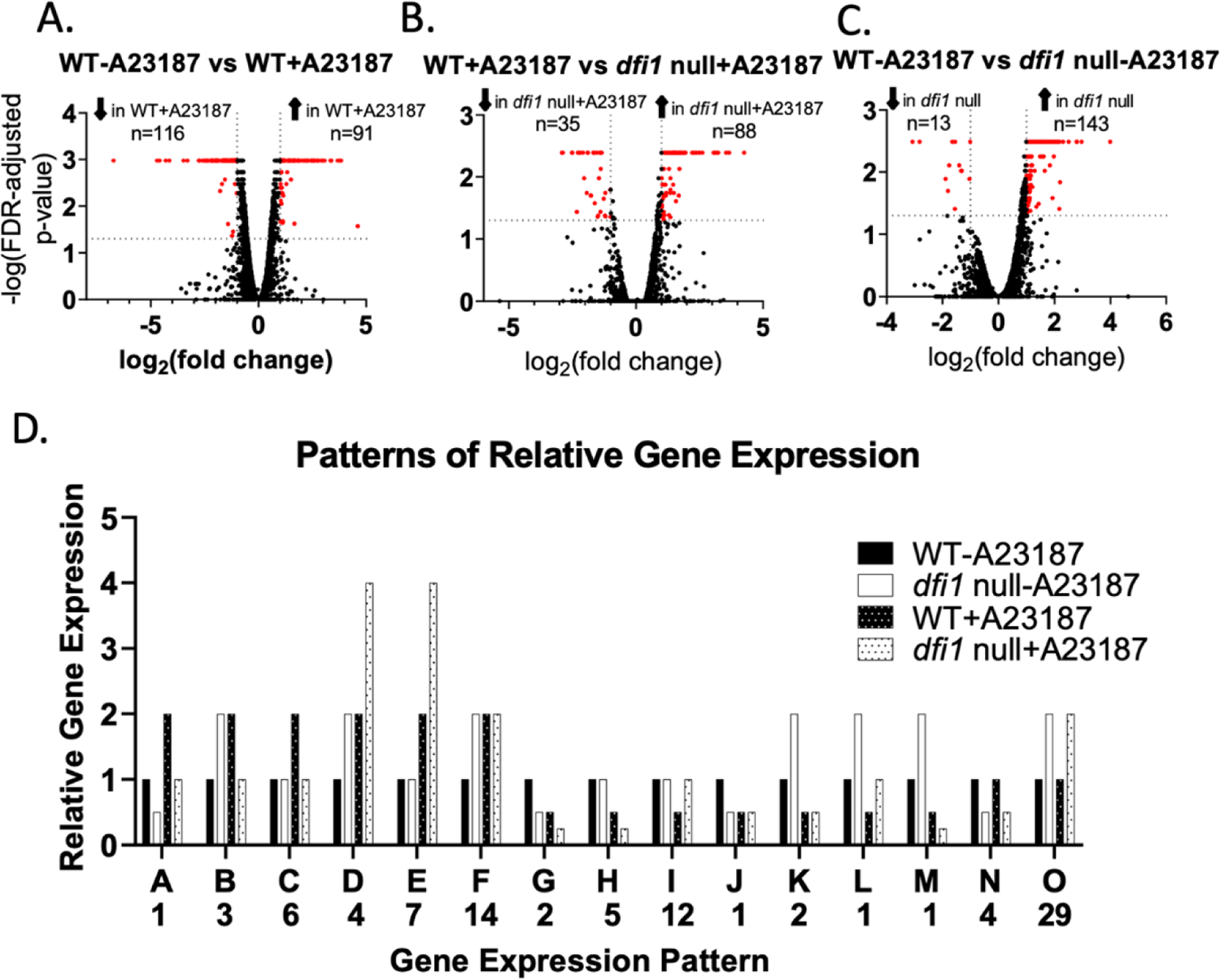
RNA-seq Identified Dfi1 Pathway-dependent gene expression. Following overnight growth in CM-U media at 25°C, WT and *dfi1* null mutant cells were treated with either the calcium ionophore A23187 (4uM) or a vehicle control (100% ethanol). After 30 minutes of treatment, cells were harvested and frozen in RNALater. RNA extracts were sent for RNA-seq analysis. Results are displayed in volcano plots. Genes in red are differentially regulated 2-fold or greater with p<0.05. The number of genes in red is displayed above plot. (A) Genes differentially expressed in WT treated with A23187 versus WT cells treated with vehicle control. (B) Genes differentially expressed in WT cells treated with A23187 vs *dfi1* null cells treated with A23187. (C) Genes differentially expressed in WT cells treated with vehicle control vs. *dfi1* null mutant cells treated with vehicle control. (D) Patterns of relative gene expression represented in the RNA-seq data. Letter below labels each pattern with corresponding information in Table 1. Number below each letter denotes the number of genes that exhibit each pattern.

**Table 1:** Patterns of Gene Expression in Response to Dfi1 Pathway Activation by A23187 Treatment

| <sup>a</sup><br>Pattern | <sup>b</sup> Description | # of<br>Genes | <sup>c</sup> Genes |
| --- | --- | --- | --- |
| **A** | Expressed at a higher level in WT vs. *dfi1* null. Expression is up-regulated by Dfi1 activation in both WT and *dfi1* null strains, but expression in activated WT cells is higher than in *dfi1* null activated cells. | 1 | **CFL5** |
| **B** | Expressed at a higher level in *dfi1* null vs. WT. Expression is up-regulated by Dfi1 activation in WT cells. Lower expression in treated *dfi1* null cells compared to untreated *dfi1* null cells. | 3 | AMO1, HGT20, NUP |
| **C** | No difference in expression in untreated WT and *dfi1* null cells. Expression increased in WT cells when Dfi1 is activated, but not in *dfi1* null cells. | 6 | **BMT9, CSA1, GAP2, HAP3, OPT1, SOD4** |
| **D** | Expressed at higher levels in *dfi1* null vs WT. Expression increases during ionophore treatment, regardless of whether Dfi1 is present. Expression of these genes is higher in treated *dfi1* null cells vs treated WT cells. | 4 | DDR48, **ICL1**, orf19.2125, orf19.6816 |
| **E** | No difference in expression in untreated WT and *dfi1* null cells. Upon ionophore treatment, expression is increased relative to the untreated controls. Upon ionophore treatment, expression increases more in *dfi1* null cells than in WT cells. | 7 | ECM331, RTA2, SLP3, orf19.2048, orf19.4476, orf19.4612, orf18.711 |
| **F** | Expressed at higher levels in *dfi1* null cells than in WT cells. Upon ionophore treatment, expression increases in WT cells. Do not respond to ionophore treatment in *dfi1* null cells. | 14 | **AAT1**, AMO2, CAN2, CIP1, GCV2, QDR1, SEO1, SHM2, SNO1, SNZ1, orf19.1306, orf19.3222, orf19.3810, orf19.6017 |
| **G** | Expressed at lower levels in untreated *dfi1* null cells than their WT counterparts. Upon ionophore treatment, expression decreases in both WT and *dfi1* null cells. | 2 | DFI1, PGA26 |
| **H** | No difference in expression in untreated WT and *dfi1* null cells. Upon ionophore treatment, expression decreases in both WT and *dfi1* null cells. Decrease in expression is greater in *dfi1* null cells than WT. | 6 | ATO1, ATO2, CSR1, POL93, ZRT2, orf19.6035 |
| **I** | No difference in expression in untreated WT and *dfi1* null cells. Upon ionophore treatment, expression decreases in WT cells, but not in *dfi1* null. | 12 | ALS1, **CCC1**, **CCP1**, **CRD2**, GLX3, HEM4, **ISA1**, MCP1, NIP7, SDH2, **SEF2**, WH11 |
| **J** | Expressed at lower levels in untreated *dfi1* null cells vs. WT. Upon ionophore treatment, expression decreases in WT cells, but shows no change of expression in *dfi1* null cells. | 1 | HGT12 |
| **K** | Expressed at higher levels in untreated *dfi1* null cells vs. WT cells. Upon ionophore treatment, expression decreases in both WT and *dfi1* null cells. Decrease is such that the difference in expression between WT and *dfi1* null is no longer significant. | 2 | OSM2, orf19.2038 |
| **L** | Expressed at higher levels in untreated *dfi1* null cells vs WT cells. Upon ionophore treatment, expression decreases in both WT and *dfi1* null cells. | 1 | BRG1 |
| **M** | Expressed at higher levels in untreated *dfi1* null cells vs WT cells. Upon ionophore treatment, expression decreases in both WT and *dfi1* null cells, but expression decreases more in *dfi1* null cells than in WT cells. | 1 | orf19.6079 |
| **N** | Expressed at lower levels in *dfi1* null vs WT. Show no response to ionophore treatment in either WT or *dfi1* null cells. | 4 | FAV1, GUT1, PGA31, orf19.938 |
| **O** | Expressed at lower levels in WT vs. *dfi1* null cells. No response to ionophore treatment in either WT or *dfi1* null cells. | 29 | ALS2, ALS4, ASR1, ASR2, BMT4, CIRT48, CSH1, FGR17, GRP2, HSP12, PEX7, STF2, UCF1, orf19.1862, orf19.2371, orf19.2959.1, orf19.33, orf19.3439, orf19.4216, orf19.5468, orf19.5514, orf19.5626, orf19.5752, orf19.6311, orf19.670.2, orf19.7085, orf19.7310 |

<sup>a</sup>Patterns of gene expression identified in Figure 1D. Gene patterns are identified by letters A-O.
<sup>b</sup>A description of the pattern of gene expression and the number of genes in each category is provided.
<sup>c</sup>The common names of each gene are listed. Gene names highlighted in bold represent genes that are members of the Sef1 regulon.

Interestingly, of these 93 genes, 13 were members of the Sef1 regulon. These genes are represented in bold in table 1. Sef1 is a transcription factor that is responsible for upregulating genes for iron uptake and iron utilization. It is active in low iron environments and its expression and activation are repressed in high iron. The Sef1 regulon is made up of 92 genes, so to have a number of these genes identified as potential targets of the Dfi1 pathway was of particular intrigue.

### Sef1 is Not Necessary for Contact-Dependent Filamentation

Based on the RNAseq data that indicated a potential connection between Dfi1, contact-dependent filamentation, and the transcription factor Sef1, a role for Sef1 in contact dependent filamentation was tested. A *sef1* null mutant strain (18) was shown to exhibit normal yeast morphology during growth in liquid culture (figure 2A, top) and did not have a growth defect in rich, high-iron media (figure 2B) or in minimal low-iron media (supplemental figure S1A-B).

**Figure 2:**
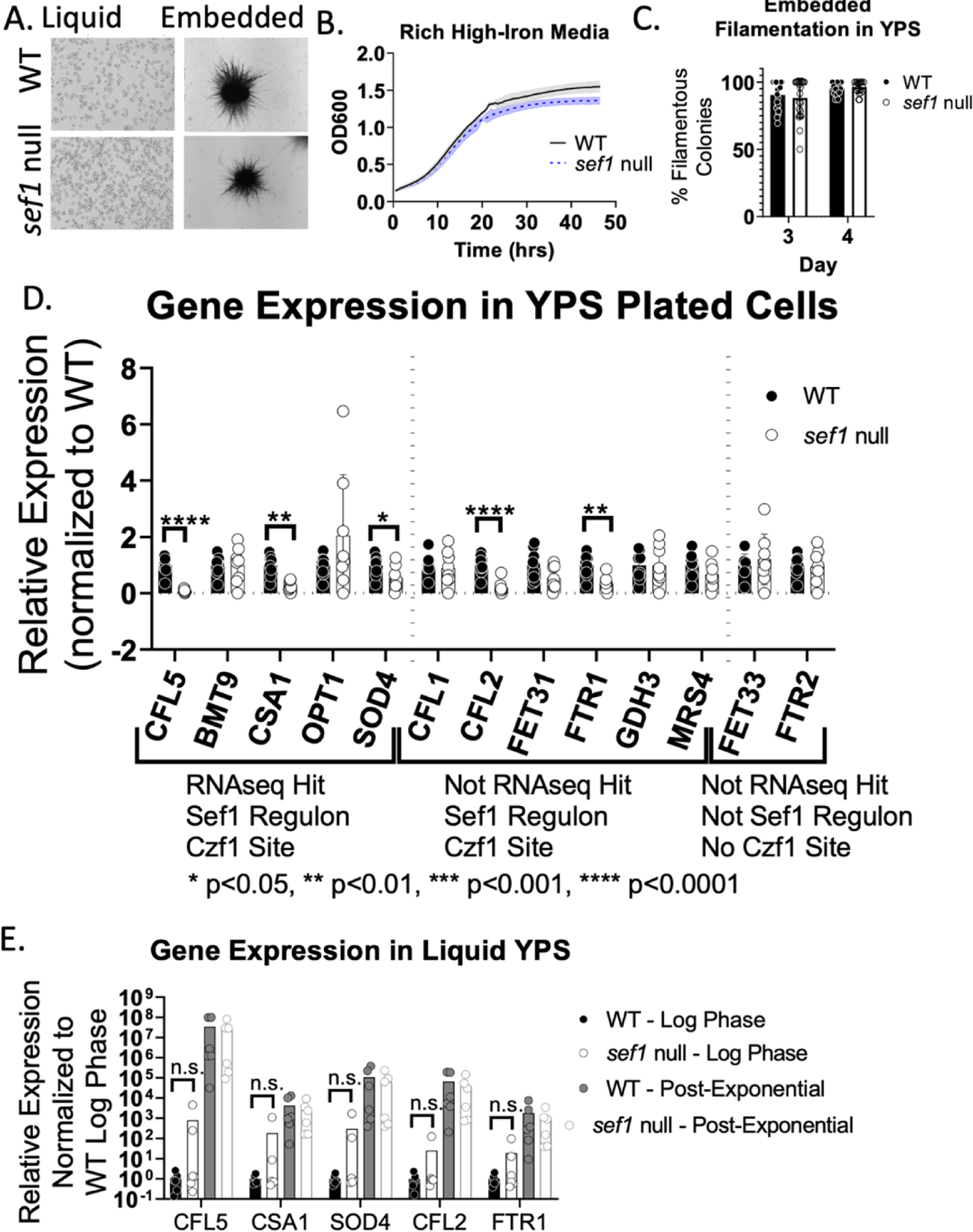
Sef1 is Not Required for Contact-Dependent Filamentation WT and *sef1* null cells were grown overnight in YPD medium at 30°C, then either back diluted to an OD of 0.1 in liquid YPS media and grown overnight at 30°C, or embedded in YPS media and allowed to grow for 4 days at 25°C, or plated on the surface of YPS plates and allowed to grow for 4 days at 25°C. (A) Brightfield images (4x objective) of WT and *sef1* null cells grown in liquid YPS (rich high-iron) medium (left) or grown embedded in YPS (rich high-iron) agar medium for 4 days at 25°C (right). (B) Growth curve of WT and *sef1* null cells in YPS media at 25°C. Black solid line, WT; gray shading, SD. Blue dotted line, *sef1* null; blue shading, SD. (C) Filamentation of WT and *sef1* null colonies grown embedded in YPS media on days 3 and 4 post-embedding at 25°C. Each point represents 1 biological replicate. 3 experiments with 3 biological replicates per experiment are shown. Bar shows the mean; error bars indicate SD (D) Gene expression in WT and *sef1* null cells grown plated on the surface of YPS media for 4 days at 25°C. Genes are labeled to indicate whether they belong to the Sef1 regulon, whether they were identified in the RNAseq study described above, and whether they contain a putative Czf1p binding site in their promoter region. Each point represents 1 biological replicate. 3 experiments with 3 biological replicates per experiment are shown. Bar shows the mean; error bars indicate SD * p<0.05, ** p<0.01, *** p<0.001, **** p<0.0001, t-tests (E) Gene expression in WT and *sef1* null cells grown to log and post-exponential phase in liquid YPS media at 25°C. Each point represents 1 biological replicate. 3 experiments with 3 biological replicates per experiment are shown. Bar shows the mean. Results are normalized to average WT expression for each experiment. Two-way ANOVA with post-hoc Dunnett’s multiple comparisons test.

To determine whether Sef1 plays a necessary role in contact-dependent filamentation, we measured the ability of the *sef1* null mutant strain to produce filamentous colonies under embedded conditions. WT and *sef1* null strains were grown embedded in YPS media as described in the Materials and Methods. Representative images of embedded colonies are shown in (figure 2A, bottom). Results from the embedded filamentation assay are shown in figure 2C. On day 3, both WT (closed symbols) and *sef1* null (open symbols) strains exhibited about 90% filamentous colonies (figure 2C, left). By day 4, both strains showed 100% filamentous colonies, and thus, no difference between WT and *sef1* null cells (figure 2C, right) was detected. This finding indicates that *SEF1* is not necessary for contact-dependent filamentation. Because expression and activation of Sef1 are repressed in high iron conditions, this assay was also performed using minimal, low-iron media. Under these conditions, a filamentation defect was not observed (supplemental figure S1C). This data indicates that Sef1 is not required for contact-dependent filamentation, in either medium condition.

To determine whether Sef1 plays a role in regulation of gene expression during growth on the surface of agar, we analyzed transcript levels for several genes. As stated above, 13 of 92 genes belonging to the Sef1 regulon were identified as potential targets of the Dfi1 pathway.

We investigated whether the expression of these genes differed from that of other Sef1 regulon genes that were not identified by the RNA-seq experiment. Additionally, we analyzed the expression of other genes that are known to be activated in low iron conditions but are not regulated by Sef1. For this analysis, we identified a collection of 13 genes to examine. The genes fall into 3 general categories – RNAseq hits that are members of the Sef1 regulon (*CFL5, BMT9, CSA1, OPT1, SOD4)*, members of the Sef1 regulon that were not RNAseq hits *(CFL1, CFL2, FET31, FTR1, GDH3, MRS4)*, and genes that are regulated by low iron (19, 20) but were not identified by RNAseq and are not members of the Sef1 regulon (*FET33, FTR2*). To investigate whether expression of these genes was altered in the *sef1* null mutant, we harvested cells grown on the surface of YPS agar medium, as described in the Materials and Methods. After harvesting the cells from the surface of the plates, the presence of invading cells was used to demonstrate the invasiveness of the strain during growth on the agar. Visual inspection using 2.5x and 10x objectives of the invading cells let behind after removing the colonies from the plates showed no discernable difference in invasive filamentation between WT and *sef1* null mutant strains (supplemental figure S1D). RNA was extracted from WT and *sef1* null mutant cells grown on YPS agar medium as described in Materials and Methods, and gene expression was examined via RT-qPCR. Despite there being no difference in filamentation, a significant decrease in expression was observed for 5 of the genes examined – *CFL5, CSA1, SOD4, CFL2,* and *FTR1* (Figure 2D).

Interestingly, the defect in expression of these 5 genes was not observed in cells grown in liquid medium. WT and *sef1* null cells grown in liquid, rich high-iron media were harvested during either log phase or post-exponential phase (4 days at 25°C). In these conditions, we observed no difference in expression of *CFL5, CSA1, SOD4, CFL2,* or *FTR1* between WT and the *sef1* null mutant (Figure 2E). These findings indicate a contact-dependent function of Sef1 that is uncoupled from the formation of filaments.

If Sef1 acts downstream of Dfi1 in the Dfi1 pathway, it is possible that there are other additional factors that also act downstream of Dfi1. Redundancy with other factors may explain why Sef1 is not necessary for filamentation under embedded conditions. A candidate factor that may also act downstream of Dfi1 is the transcription factor Czf1.

Czf1 is a zinc-cluster DNA binding protein that plays a role in contact-dependent filamentation (21, 22). Recently, Czf1 has been shown to be a regulator of cell wall architecture and integrity and is also required for basal levels of caspofungin tolerance (23). Embedding a *czf1* null strain as described above in rich high-iron media showed a defect of filamentation at early time points (figure 3A-B), consistent with previously published data (24). Of the 92 genes in the Sef1 regulon, 74 (80%) contain a putative Czf1p binding site (TTWRSCGCCG (25)) in their promoter (defined here as the entire upstream intergenic region). To compare this to the prevalence of the Czf1p binding site in the *C. albicans* genome overall, 200 genes were randomly selected and their promoter regions (entire upstream intergenic region) scanned for Czf1p binding sites. Of these 200 randomly selected genes, only 27% contained a Czf1p binding site in their promoter. This represents a significant enrichment of Czf1 binding sites among Sef1 regulon genes (p<0.0001, Fisher’s exact test), leading us to hypothesize that Czf1 may regulate genes in the Sef1 regulon. Analysis of transcripts by RT qPCR of the same 13 genes listed above from WT and *czf1* null cells plated on rich high-iron media showed a significant decrease in expression of 9 genes – *CFL5, BMT9, CSA1, OPT1, SOD4, FET31, FTR1, GDH3, MRS4,* and *FET33* (Figure 3C). Interestingly, 4 of the 5 genes differentially regulated in the *sef1* null strain also require Czf1 for WT levels of expression; only *CFL2* required Sef1 but not Czf1. Furthermore, all of the genes in this collection that were identified by the RNAseq experiment as potential targets of the Dfi1 pathway required Czf1 for WT levels of expression in contact conditions. Therefore, we have identified genes whose expression requires Sef1 only (*CFL2)*, Czf1 only (*BMT9, OPT1, FET31, GDH3, MRS4,* and *FET33*), or both Sef1 and Czf1 (*CFL5, CSA1, SOD4,* and *FTR1)* for WT levels of expression during growth in contact conditions.

**Figure 3:**
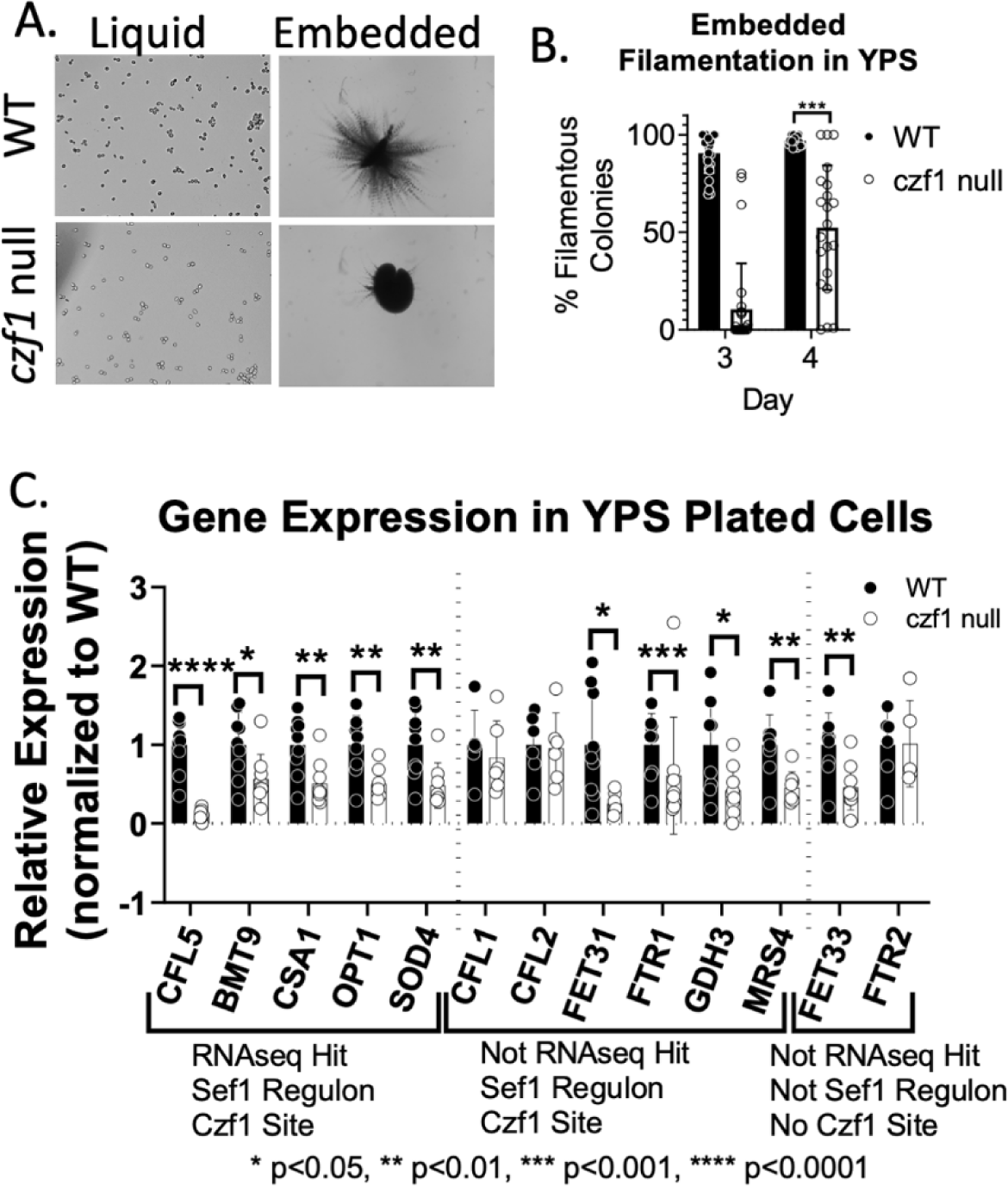
Czf1 is Required for Gene Expression in Plated Cells WT and *czf1* null cells were grown overnight in YPD medium at 30°C, then either back diluted to an OD of 0.1 in YPS media and grown overnight at 30°C, embedded in YPS media and allowed to grow for 4 days at 25°C, or plated on the surface of YPS plates and allowed to grow for 4 days at 25°C. (A) Brightfield images (4x objective) of WT and *czf1* null cells grown in liquid or grown embedded in YPS (rich high-iron) media. (B) Filamentation of WT and *czf1* null cells grown embedded in YPS media on days 3 and 4 post-embedding. (C) Gene expression in WT and *czf1* null cells grown plated on YPS media for 4 days. Genes are labeled to indicate whether they belong to the Sef1 regulon, whether they were identified in the RNAseq study described above, and whether they contain a putative Czf1p binding site in their promoter. Results are normalized to average WT expression for each experiment. (B & C) Each point represents 1 biological replicate. 3 experiments with 3 biological replicates per experiment are shown. Bar shows mean; error bars show SD. * p<0.05, ** p<0.01, *** p<0.001, **** p<0.0001, t-tests

In summary, Sef1 is not required for contact-dependent filamentation, while WT levels of embedded filamentation require Czf1. Both transcription factors are required for WT levels of *CFL5, CSA1, SOD4* and *FTR1* expression in cells growing on the surface of agar. However, despite 5 genes demonstrating Sef1-dependent expression during plated growth, none of these genes required Sef1 for WT levels of expression during liquid growth, indicating a potential role for Sef1 in contact-dependent filamentation.

### Activated Sef1 is Sufficient to Overcome the *dfi1* Null Filamentation Defect

To determine whether Sef1 could play a functional role in the Dfi1 pathway and affect contact-dependent filamentation, we asked whether constitutive activation and expression of Sef1 would be sufficient to overcome the filamentation defect of a *dfi1* null mutant. An activated Sef1 fusion (kindly provided by Dr. Joachim Morschhäuser, University of Würzburg) was used. The activated allele encodes a fusion of a Gal4 activation domain (GAD) to the C-terminus of Sef1, causing the protein to be constitutively activated. The activated *SEF1* allele is constitutively expressed under control of the *ADH1* promoter (26). This construct was transformed into WT and *dfi1* null strains by electroporation and transformants were selected as described in Materials and Methods. All transformants were confirmed by PCR. The strains did not exhibit aberrant morphology when grown in liquid rich high-iron media (Figure 4A).

**Figure 4:**
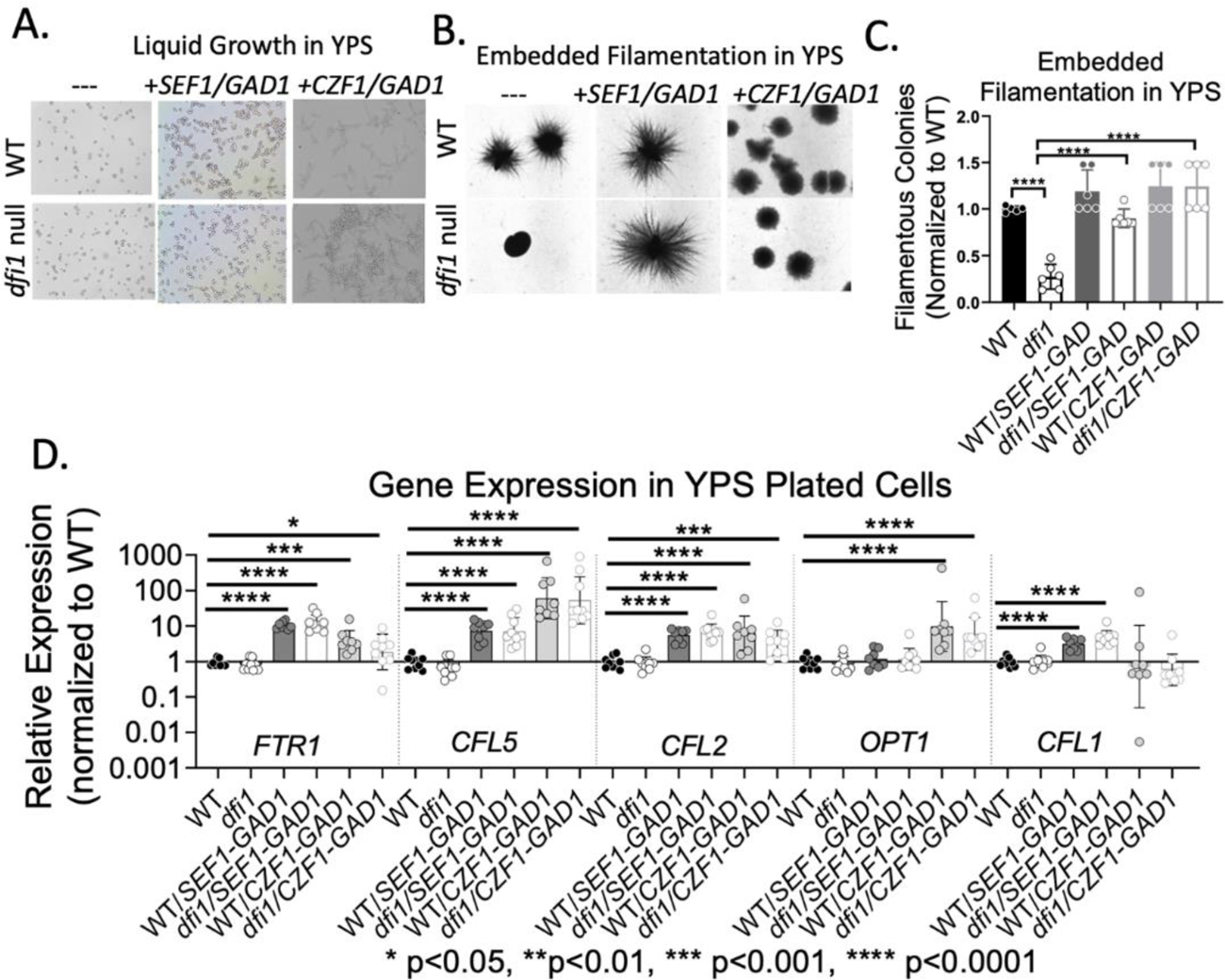
Constitutive Activation of Sef1 is Sufficient to Overcome *dfi1* Null Contact-Dependent Filamentation Defect WT, *dfi1* null, WT+*SEF1/GAD*, *dfi1+SEF1/GAD*, WT+*CZF1/GAD*, and *dfi1*+*CZF1/GAD* cells were grown overnight in YPD medium at 30°C, then either back diluted to an OD of 0.1 in YPS media and grown overnight at 30°C, embedded in YPS media and allowed to grow for 4 days at 25°C, or plated on the surface of YPS plates and allowed to grow for 4 days at 25°C. (A) Brightfield images (4x objective) of WT and *dfi1* strains alone, with *SEF1/GAD*, or with *CZF1/GAD* alleles grown in liquid YPS. (B) Brightfield images (4x objective) of WT and *dfi1* strains alone, with *SEF1/GAD*, or with *CZF1/GAD* alleles grown embedded in YPS. (C) Relative number of filamentous colonies for WT, *dfi1*, WT+*SEF1/GAD, dfi1+SEF1/GAD,* WT+*CZF1/GAD,* and *dfi1*+*CZF1/GAD* strains grown embedded in YPS media. Each point represents 1 biological replicate. 3 experiments with 3 biological replicates per experiment are shown. Results were normalized to average WT % filamentous colonies for each experiment. (D) Relative gene expression in WT, *dfi1*, WT+*SEF1/GAD, dfi1+SEF1/GAD,* WT+*CZF1/GAD,* and *dfi1*+*CZF1/GAD* strains grown on the surface of YPS plates. Each point represents 1 biological replicate. 3 experiments with 3 biological replicates per experiment are shown. Results are normalized to average WT expression for each experiment. * p<0.05, ** p<0.01, *** p<0.001, **** p<0.0001, two-way ANOVA with post-hoc Dunnett’s multiple comparisons test

WT, *dfi1*, WT/*SEF1-GAD*, and *dfi1/SEF1-GAD* strains were embedded in rich, high-iron media as described in Material and Methods. On day 4, the WT strain exhibited filamentous colonies. The *dfi1* null mutant strain yielded only 25% as many filamentous colonies as the WT (p<0.0001; one-way ANOVA with post-hoc Tukey multiple comparisons test), confirming the filamentation defect for the *dfi1* mutant that has been previously reported (12) (Figure 4B-C, supplemental figure S2). The WT/*SEF-GAD* strain exhibited filamentation consistent with WT, indicating that constitutive activation of Sef1p did not increase filamentation (Figure 4B, supplemental figure S2). When the *SEF1-GAD* allele was added to the *dfi1* null strain, 100% of the scored colonies exhibited filamentation comparable to WT, resulting in a statistically significant difference in filamentous growth between the *dfi* null strain and the *dfi1/SEF1-GAD* strain (p=<0.0001; one-way ANOVA with post-hoc Tukey multiple comparisons test). Activated Sef1p was thus sufficient to rescue the filamentation defect seen in the *dfi1* null mutant strain (figure 4C). Additional WT and *dfi1* null strains transformed at the same locus with a *SAT1* cassette not encoding the *SEF1-GAD* allele were also characterized and showed no rescue of the *dfi1* null filamentation defect (data not shown). Thus, activation of Sef1 was sufficient to bypass the filamentation defect caused by lack of Dfi1. Sef1 activation, however, did not lead to filamentation under liquid growth conditions.

Embedded filamentation by these strains was also analyzed at 37°C. Under these conditions, the *dfi1* null mutant did not exhibit a consistent defect in filamentation and the *SEF1-GAD* allele did not increase filamentation. Most likely, alternative filamentation regulatory pathways are activated at 37°C and these pathways mask the defect of the *dfi1* null mutant and prevent the detection of bypass by the *SEF1-GAD* allele (data not shown).

To examine how constitutive expression and activation of Sef1 affects gene expression, the same cultures that were embedded were also plated on the surface of YPS agar plates, as described above. Cells were harvested from the agar after 4 days of growth at 25°C and RNA was extracted as described in Materials and Methods. Five genes were selected for analysis – *FTR1, CFL5, CFL2, OPT1,* and *CFL1.* These genes were selected because of their varying dependence on Sef1 and Czf1 as shown in Figs. 2D and 3C. These genes are all members of the Sef1 regulon, however only *CFL5, FTR1,* and *CFL2* required Sef1 for WT levels of expression in plated cells grown on rich, high-iron media. We observed that the WT/*SEF1-GAD* strain exhibited a 10-fold increase in expression of *FTR1, CFL5, CFL2,* and *CFL1* relative to WT (Figure 4D). The *dfi1/SEF1-GAD* strain exhibited expression of *FTR1, CFL5, CFL2,* and *CFL1* similar levels to its WT counterpart (Figure 4D). In contrast, constitutive expression and activation of Sef1 did not appear increase *OPT1* expression under these growth conditions, consistent with the data shown in Figure 2D. Together, these results indicate that while Sef1 was not necessary for contact-dependent filamentation, its activation was sufficient to bypass Dfi1 and promote embedded filamentation. Additionally, constitutive activation of Sef1 led to expression of some genes beyond their WT levels of expression.

When a similar constitutively expressed and activated *CZF1* allele was introduced into WT and *dfi1* null strains, the cells exhibited enhanced filamentation. Both WT/*CZF1-GAD* and *dfi1/CZF1-GAD* strains formed filaments when grown in liquid culture (figure 4A). Embedding these strains as described above resulted in highly filamentous colonies with filaments that appeared to differ from their WT counterparts. As shown in figure 3B, strains containing the *CZF1-GAD* allele exhibited shorter filaments, compared to the longer filaments of the WT strain.

As with Sef1, we observed a rescue of the *dfi1* null filamentation defect due to constitutive expression and activation of Czf1 (p<0.0001) (Figure 4C). Similar to above, we analyzed gene expression in WT/*CZF1-GAD* and *dfi1/CZF1-GAD* strains grown on the surface of YPD agar. All of the genes analyzed contained putative Czf1 binding sites in their promoters (as defined above), but only *FTR1, CFL5,* and *OPT1* required Czf1 for WT levels of expression in plated conditions (figure 3C). Analysis of gene expression in plated cells showed an increase in expression in 4 of the 5 genes (*FTR1, CFL5, CFL2,* and *OPT1*) when constitutively expressed and activated Czf1 was present (figure 4D). Constitutive activation and expression of Czf1 was not sufficient to induce higher levels of expression of *CFL1.* These results are consistent with the observation that *CFL5, FTR1,* and *OPT1* required Czf1 for WT levels of expression but *CFL1* did not. Constitutive activation of Czf1 was also not sufficient to raise expression of *CFL1* over WT levels.

Interestingly, a recent study showed that constitutive expression and activation of Czf1 was also sufficient to increase expression of *CFL5* and *OPT1* during growth in liquid in a liquid growth model (23). Taken together, these data showed that both Sef1 and Czf1 are capable of regulating invasive filamentation but demonstrate a complex pattern of gene expression dependent on the factors available and contact conditions.

### Low Iron conditions increased Contact-Dependent Filamentation by a *dfi1* Null Mutant

Physiologically, Sef1 is activated during growth in low-iron conditions, and potentially its activation under these conditions could bypass the filamentation defect observed in the *dfi1* mutant. To test this hypothesis, we grew cells in YNB media containing different levels of iron (adapted from Hsu et al (27)). Cells were embedded in this medium as described in Materials and Methods. On day 4, colonies were inspected for evidence of invasive filamentation. As expected, WT colonies were nearly 100% filamentous in high iron (50-500 uM) conditions (figure 5A, B), consistent with filamentation in rich, high-iron media (12). As the amount of iron in the media was decreased, WT cells retained their ability to filament identically to their high-iron counterparts. It was only in the absence of any added iron that WT colonies were non-filamentous (figure 5B, 0 uM). These colonies were also smaller than those in higher iron conditions, indicating that the lack of iron inhibited normal growth.

**Figure 5:**
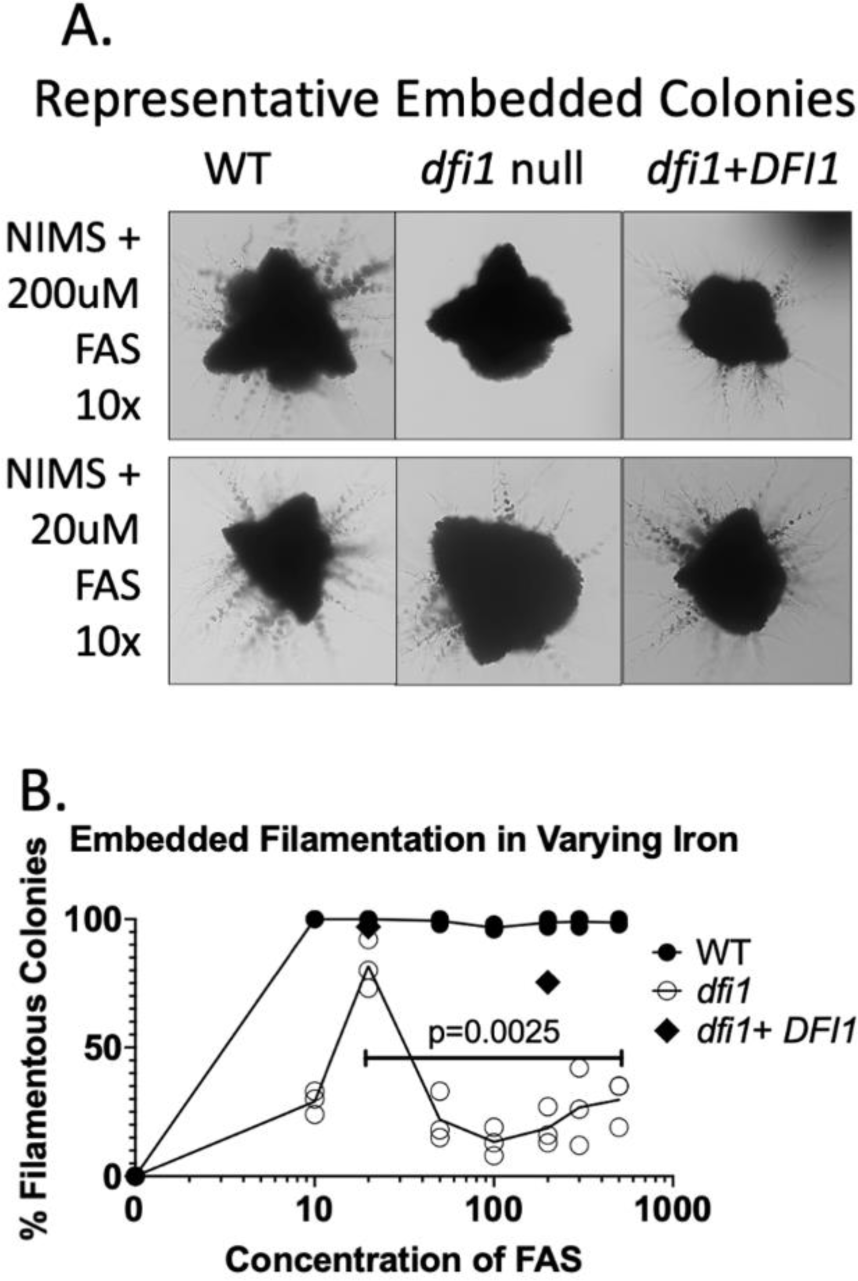
Low-Iron Medium Conditions increase Contact-Dependent Filamentation in the *dfi1* null mutant. WT and *dfi1* null cells were grown overnight in YPD medium at 30°C, then back diluted to an OD of 0.1 in NIMS media supplemented with 200 uM or 20uM FAS and grown overnight at 30°C or embedded in the same media and allowed to grow for 4 days at 25°C. (A) Brightfield images (10x objective) of WT, *dfi1* null, and *dfi1*+*DFI1* cells grown embedded in minimal high-iron or minimal low-iron media. (B) Filamentation of WT, *dfi1* null, and *dfi1*+*DFI1* cells grown in iron-free media supplemented with a range of concentrations of Ferrous Ammonium Sulfate (FAS). Each point represents the average of 1 experiment with 3 biological replicates; 3 experiments are shown. ** p<0.01, t-test

In relatively high concentrations of iron (50-500 uM), we observed that the *dfi1* mutant exhibited lower levels of filamentous colonies (figure 5A, B) consistent with the defect in filamentation observed in rich, high-iron medium. However, at 20 uM iron, we observed an increase in the number of colonies that were scored as filamentous (figure 5B), indicating that low-iron conditions led to a partial bypass of the *dfi1* null mutant defect in filamentation in embedded conditions. This rescue of filamentation was limited to contact conditions, as growth in liquid medium at 20 uM iron did not result in filamentation (supplemental figure S3). In very low concentrations of iron (0-10 uM), *dfi1* null mutant colonies did not exhibit filamentation (figure 5B). Upon closer examination, these colonies were also much smaller than their higher iron counterparts, indicating that the lack of available iron was inhibiting growth. Consistent with previous results, introduction of a WT allele of *DFI1* into the *dfi1* null mutants restored filamentation during growth under embedded conditions (Figure 5B, diamond symbols, 20uM and 200uM FAS). Since Sef1 is induced in low iron conditions, our model is that this filamentation recovery is due to induction and activation of Sef1p due to low iron. These results demonstrate an effect of Sef1 on filamentation during growth in contact with agar medium.

## Discussion

Dfi1 is a plasma membrane protein that activates an embedded filamentation signaling pathway. The Dfi1 signaling pathway is required for WT levels of invasive filamentation in colonies grown under embedded conditions at 25°C. At 37°C, the *dfi1* mutant does not exhibit a defect in filamentation in embedded conditions, presumably because other filamentation signaling pathways are also active. Here we identified two transcription factors that are effectors of the Dfi1 signaling pathway – Czf1 and Sef1. Czf1, a zinc cluster DNA binding protein, was previously shown to be required for WT filamentation under embedded conditions at low temperature but not under other conditions. The work described here showed that Czf1 is required for expression of several genes in cells growing on the surface of agar (figure 3C).

Constitutive activation of Czf1 promoted embedded filamentation in the absence of Dfi1 and but was not sufficient to increase gene expression of the analyzed genes over their WT levels. These results support the model that Czf1 functions downstream of Dfi1 in a pathway that regulates embedded filamentation.

Less expectedly, we identified a role for low iron and the zinc cluster DNA binding protein Sef1 in Dfi1-mediated contact-dependent filamentation. While *SEF1* is not necessary for contact-dependent filamentation, it is necessary for expression of a number of genes (*CFL5, CSA1, CFL2,* and *FTR1*) in contact conditions (figure 2C). Constitutive expression and activation of *SEF1* resulted in a rescue of the *dfi1* null contact-dependent filamentation defect, demonstrating that activated Sef1 can influence filamentation and was also sufficient to increase gene expression in a number of genes. Additionally, we demonstrated that decreasing the amount of iron present in the media resulted in a partial rescue of the *dfi1* null contact-dependent filamentation defect, again consistent with the notion that activation of Sef1 increased filamentation. This evidence indicates a role for both Sef1 and Czf1 in Dfi1-mediated contact-dependent filamentation.

Czf1 and Sef1 are both required for WT levels of expression of *CFL5* and *FTR1* in plated cells. These results support a cooperative model of *CFL5* and *FTR1* regulation by Sef1 and Czf1 in which the two factors function together to promote gene expression under these conditions. The putative binding site for Czf1 is present in the promoters of an estimated 27% of genes in the *C. albicans* genome and in the promoters of about 80% of the genes belonging to the Sef1 regulon, including *CFL5, FTR1,* and other genes analyzed. Thus, there may be substantial overlap between the Sef1 and Czf1 regulons. Activated Sef1 may be able to activate expression of genes that are usually regulated by Czf1, and promote embedded filamentation under certain conditions.

Based on the evidence provided above, Sef1 and Czf1 have different effects on filamentation and gene expression. *C. albicans* requires Czf1 but not Sef1 for normal contact-dependent filamentation in YPS agar medium at low temperature. Sef1 may have a backup function in regulating contact-dependent filamentation under these conditions but could have a more prominent role under other conditions. Because expression and activation of Sef1 is normally repressed by Sfu1 in high-iron conditions, it is conceivable that there is not enough Sef1 present in YPS agar growth conditions to compensate for the lack of Czf1. Interestingly though, Sef1 is still required for expression of a number of genes in high iron conditions and thus, the low levels of active Sef1 that are present may function together with Czf1 to bring about normal gene expression.

Taken together, we hypothesize the following model for Dfi1, Sef1, and Czf1 interactions (figure 6): In normal, relatively high iron media, when Dfi1 is present, activation of Dfi1 in response to a contact signal results in Czf1 activation. Activated Czf1 binds to promoters and allows activation of gene expression leading to filamentation. Under these conditions, the Dfi1 pathway also activates Sef1, but Sef1 is not required for filamentation because Czf1 is present. In the absence of Dfi1, Czf1 and Sef1 are not activated and embedded filamentation is defective. However, in low iron media, low iron availability can trigger expression and activation of Sef1 and Sef1 expression and activation increases filamentation of the *dfi1* mutant.

**Figure 6:**
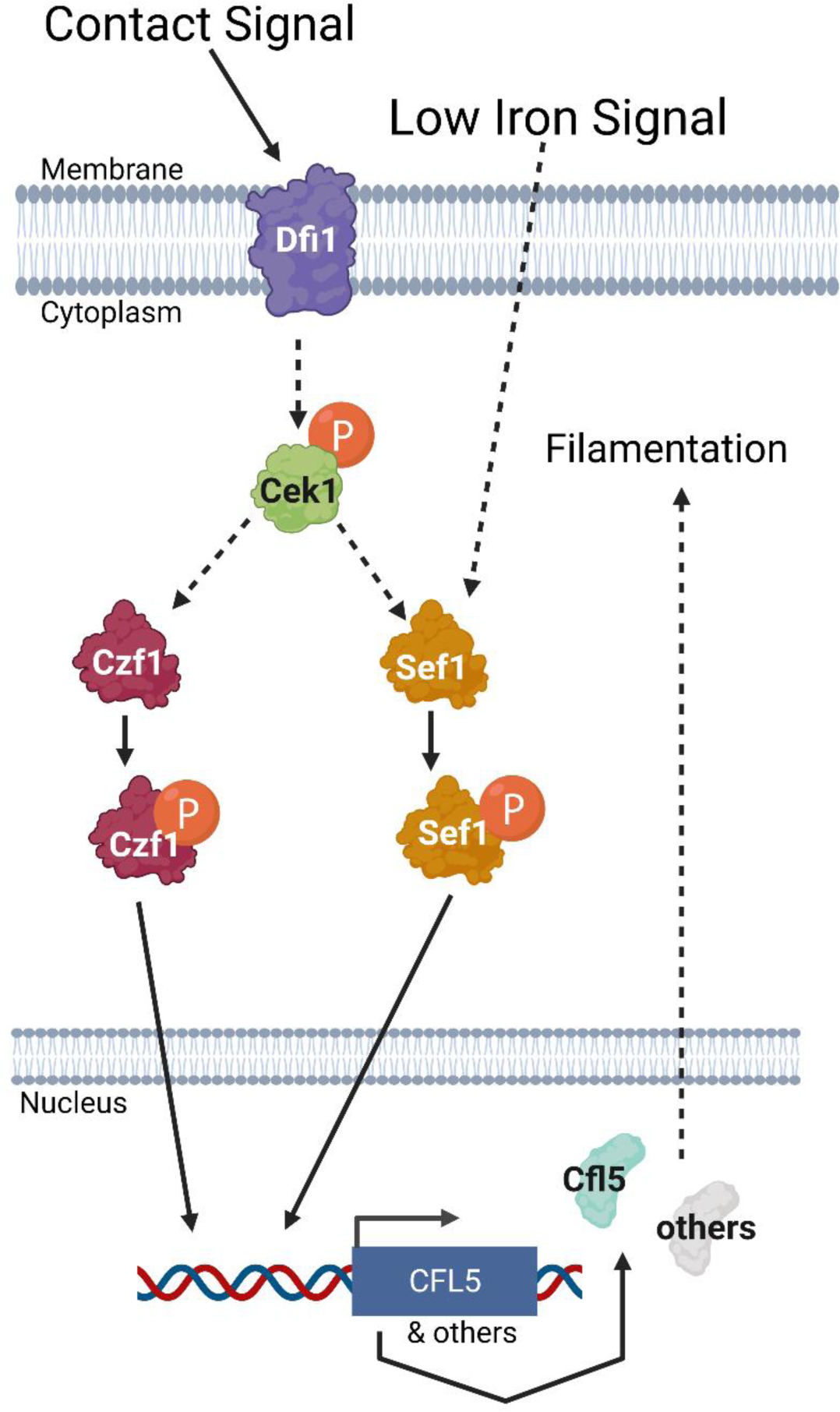
Proposed Model of Dfi1, Sef1, and Czf1 During Contact-Dependent Filamentation The hypothesized model for interactions between Dfi1, Sef1, and Czf1 is as follows: When cells respond to growth in contact with an agar medium, signaling proceeds through Dfi1, resulting in downstream Cek1 activation. Cek1 activation leads to activation of Czf1, resulting in translocation into the nucleus and subsequent gene expression leading to filamentation. Cek1 activation through Dfi1 can also lead to Sef1 activation and downstream gene expression, but this is not required when Czf1 is present. Sef1 can also be activated in low iron media, and in the absence of Dfi1 (and therefore Czf1 activation), this Sef1 activation is sufficient to result in contact-dependent filamentation. Finally, constitutive activation of either Sef1 or Czf1 results in contact-dependent filamentation, even in the absence of Dfi1.

The gastrointestinal tract is an iron-replete environment (14). In the GI tract, activation of the Dfi1 pathway in response to a contact signal could lead to activation of Sef1, promoting expression of genes such as those described above, which are expressed in plated cells in a Sef1-dependent manner. If Sef1 is activated by the Dfi1 pathway coincident with the initiation of tissue invasion, the cells will be equipped to compete successfully for iron before they actually encounter the low iron environment that is characteristic of tissue. Hence early activation of Sef1 due to the action of the Dfi1 pathway may enhance the rapidity with which invading cells adapt to a change in iron availability.

Further, the fact that Sef1 also plays a role in contact-dependent filamentation highlights an intersection between filamentation and iron uptake. Other iron uptake genes, such as *CFL1*, have been shown to have functions in filamentation, with deletions leading to impaired filamentous growth and altered cell wall architecture in liquid conditions. Recently, Luo et al showed that iron acquisition was required for sustained hyphal development, but not hyphal initiation (28). Furthermore, they suggested that Sef1 can be activated in response to the same stimuli that induce hyphal growth in order to facilitate expression of iron uptake genes (28). Here, we propose that under the conditions of our experiments, hyphal inducing conditions activate Dfi1, and Dfi1 activation results in Sef1 and Czf1 activation. Further, we observed a rescue of the *dfi1* filamentation defect specific to contact conditions when Sef1 was activated. Luo et al postulated that hyphal-development is itself an iron consuming process, and therefore that the act of invading media creates an iron-poor environment, leading to the necessity of iron uptake (28). Our findings are consistent with this model.

As described above, we observed that a number of genes, including *FTR1*, are dependent on Sef1 and Czf1 for WT levels of expression in contact conditions. Previous studies by others have shown that deletion of *FTR1* leads to attenuated virulence (29), and *FTR1* transcript levels have been reported to be increased during hyphal elongation. Regulation of *FTR1* expression may contribute to the effects of Dfi1 on invasive filamentation and the ability to produce a lethal infection in the intravenously inoculated mouse (12). Thus, the interconnection between the invasive filamentation pathway and the iron uptake system mediated by Dfi1 contributes to the pathogenicity of *C. albicans*.

## Materials and Methods

### Strains and Growth Conditions

All strains used are detailed in Table 2. *C. albicans* was routinely cultured using Yeast Extract Peptone Dextrose (YPD) (1% Yeast Extract, 2% Peptone, 2% glucose) medium at 30°C. For specific studies, cells were grown in complete minimal medium minus uridine (CM-U) or Yeast Extract Peptone Sucrose (YPS) (1% Yeast Extract, 2% Peptone, 2% Sucrose). Non-Iron Media was adapted from Hsu et al (26) as follows: Yeast Nitrogen Base (YNB) minus Fe, Mn, Zn, and Cu (USBiological) was supplemented with 2.37uM MnSO4, 1.39uM ZnSO4, and 0.25uM CuSO4 to reflect their normal concentrations in YNB. Bathophenanthroline disulfonic acid (BPS) at 100 µM concentration was used to remove any residual iron. Ferrous ammonium sulphate was added at 0 – 500 uM concentrations, as well as 2% sucrose. To create embedded conditions, 0.8% agarose was added and cells were embedded as described above. Cells were routinely cultured at 30°C or 25°C. Some mutants were obtained as part of a deletion collection (18). All deletions were confirmed by PCR.

**Table 2:** List of Strains Used in This Study

| Strain | Description | Genotype | Source |
| --- | --- | --- | --- |
| pcz1 | wildtype | BWP17, *ura3Δ::imm434/URA3* | Zucchi et al, 2011 |
| pcz5 | *dfi1* null | *Pcz1, dfi1Δ/dfi1Δ*<br>*ura3Δ::imm434/URA3* | Zucchi et al 2011 |
| pcz9 | *dfi1/DFI1* | *pcz5, dfi1Δ/dfi1::DFI1-His<sub>6</sub>HA-*<br>*HIS* placer | Zucchi et al 2011 |
| SN425 | wildtype | SN152, *his1Δ/HIS1, leu2Δ/LEU2* | Homann et al, 2009 |
| TF015-Y | *sef1* null | SN152, *sef1Δ::HIS1/sef1Δ::LEU2* | Homann et al, 2009 |
| TF104-X | *czf1* null | SN152, *czf1Δ::HIS1/czf1Δ::LEU2* | Homann et al, 2009 |
| arj1 | WT+SEF1/GAD | *pcz1, ADH1/adh1::P<sub>ADH1</sub>-SEF1-*<br>*GAL4AD-HA-caSAT1* | This work |
| arj2 | *dfi1*+SEF1/GAD | *pcz5, ADH1/adh1::P<sub>ADH1</sub>-SEF1-*<br>*GAL4AD-HA-caSAT1* | This work |
| arj3 | WT+CZF1/GAD | *pcz1, ADH1/adh1::P<sub>ADH1</sub>-CZF1-*<br>*GAL4AD-HA-caSAT1* | This work |
| arj4 | *dfi1*+CZF1/GAD | *pcz5, ADH1/adh1::P<sub>ADH1</sub>-CZF1-*<br>*GAL4AD-HA-caSAT1* | This work |

### Strain Construction

*C. albicans* strains were transformed by electroporation as described in Reuss et al (30). Plasmids containing *SEF1-GAD* and *CZF1-GAD* alleles were generously provided by the Morschhauser lab. Described in Schilling and Morschhauser (26), these were integrated into the genomes of WT and *dfi1* null strains at the *ADH1* locus. The presence of a *SAT1* cassette allowed for the selection of transformants via nourseothricin resistance. Presence of the *SEF1-GAD* and *CZF1-GAD* alleles were confirmed by PCR and agarose gel electrophoresis. Four independently isolated strains of WT/*SEF1-GAD* and *dfi1/SEF1-GAD* were characterized.

### Artificial Activation of Dfi1p Pathway

Artificial activation of the Dfi1p pathway in liquid medium was done as previously described in Davis et al (13). Briefly, *C. albicans* cells were either treated with 4 uM of the calcium ionophore A23187 or 100% ethanol as a vehicle. After 30 minutes, 10 mL of cells were collected for RNA extraction and analysis. Cells for RNA were washed and frozen in RNALater at −80°C.

### RNA Extraction

RNA was extracted from *C. albicans* cells frozen in RNALater using the Qiagen RNeasy kit with the following modifications: Cells were broken by bead beating with 0.5 mm silica zirconia beads on a Minibeadbeater 24 machine (Biospec) with 3 rounds of 1 minute bead beating and 5 minutes on ice between. RNA was extracted from the bead beating supernatant as described by Qiagen, including on column DNase treatment, and eluted with 30uL of water. RNA was stored at −80°C.

### RNA Sequencing

RNA was sent to the Tufts University Core Facility for library preparation and sequencing using the Illumina TruSeq RNA library preparation kit and the HiSeq 2500 instrument.

Sequencing data was analyzed using the Tuxedo Suite as previously described (31). NCBI genome Candida albicans SC5314 (assembly ASM18296v3) (32) and annotation GFF file were used as reference genome. A bowtie2 index was created using bowtie2-build (v2.2.1) from the fna sequence file, while the downloaded GFF was converted to GTF using gffread for downstream use. The raw sequencing reads for each sample were mapped to the bowtie2 index using bowtie2 (v2.2.1) with default parameters, and then sorted and converted to bam format using samtools v1.9 (sort function). The resulting sorted bam files were used as input for cuffdiff (cufflinks v2.1.1) with the triplicates grouped. Genes with a fold change of over 2 were considered.

### Embedded Filamentation Assays

The growth of colonies under embedded conditions was performed as previously described in Zucchi et al (12). Briefly, lukewarm 1% YPS agar was pipetted onto a drop of medium containing approximately 150 cells. Three replicate cultures of each strain were independently embedded and plates were placed in a humidified chamber at 25°C for four days. After 4 days of growth, embedded colonies were microscopically examined using 4x and 10x objectives for evidence of filamentation. A colony was considered a “filamentous colony” if it contained 20 or more visible filaments. The reported “percent filamentous colonies” refers to the percentage of colonies counted that meet this criterium; 75-125 colonies were counted per plate. Plates were scored blinded to prevent counting bias. All filamentation assays were repeated 3 times.

### Growth of Strains on the Surface of Agar Medium

Cells were grown as previously described (12). Briefly, cells were plated to obtain single colonies on YPS (1% Yeast Extract, 2% Peptone, 2% sucrose) with 1% agar and grown at 25°C for four days. Cells were washed off the plate with RNALater and frozen at −80°C for later RNA analysis.

### RT-qPCR

cDNA was synthesized by reverse transcription of 10ug of total RNA using SSIII (Invitrogen) following manufacturer’s protocol. Resulting cDNA was diluted 1:20 and used for gene expression analysis via qPCR. qPCR reactions were set up using SYBR Green MasterMix (AppliedBiosystems). qPCR reactions were run on AppliedBiosystems StepOnePus RT-qPCR system using standard reaction parameters. Primers used for qPCR are listed in (Table 3). All qPCR products were confirmed via sequencing. Samples with no template or with RNA that was not converted to cDNA did not yield products.

**Table 3:**
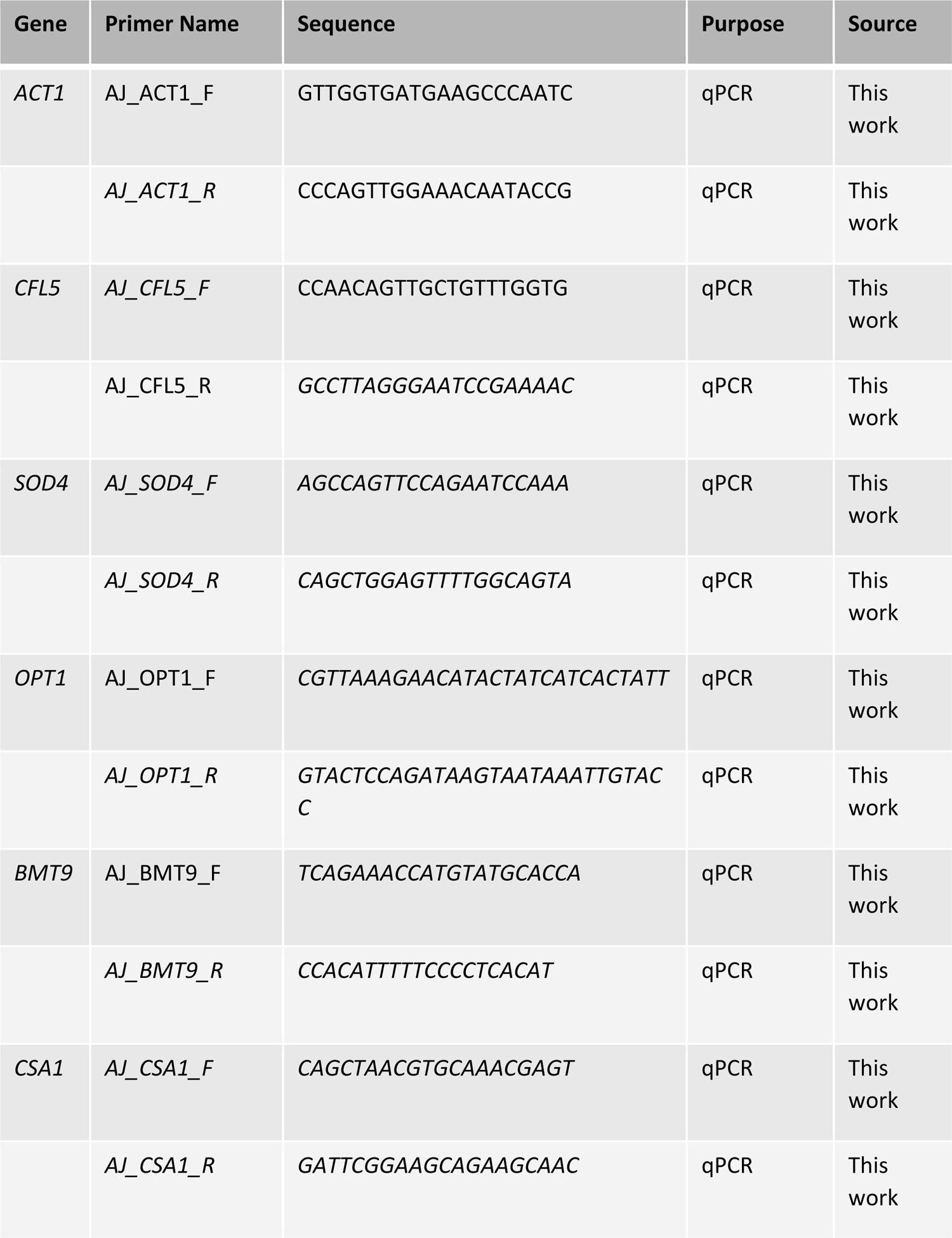
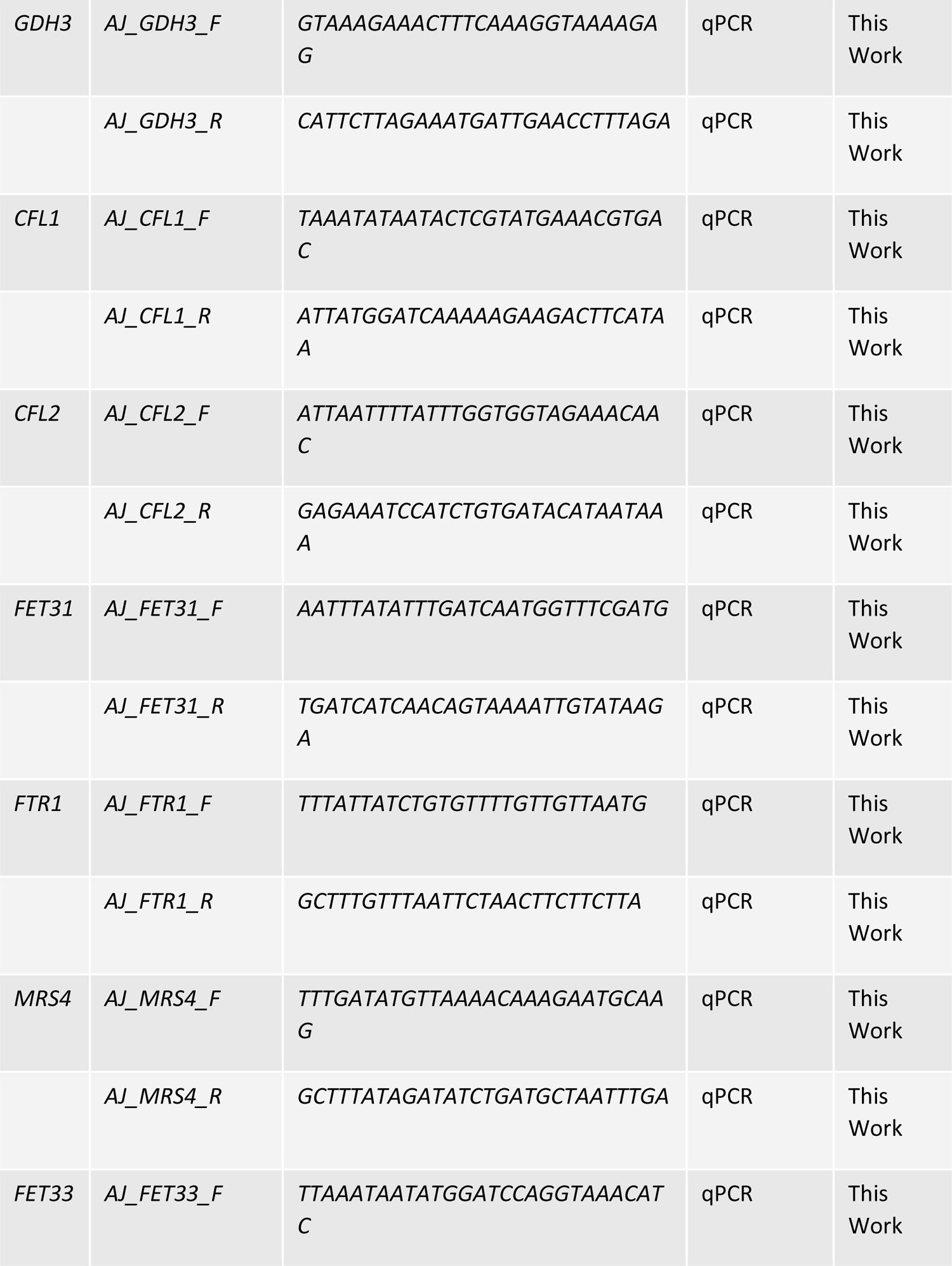
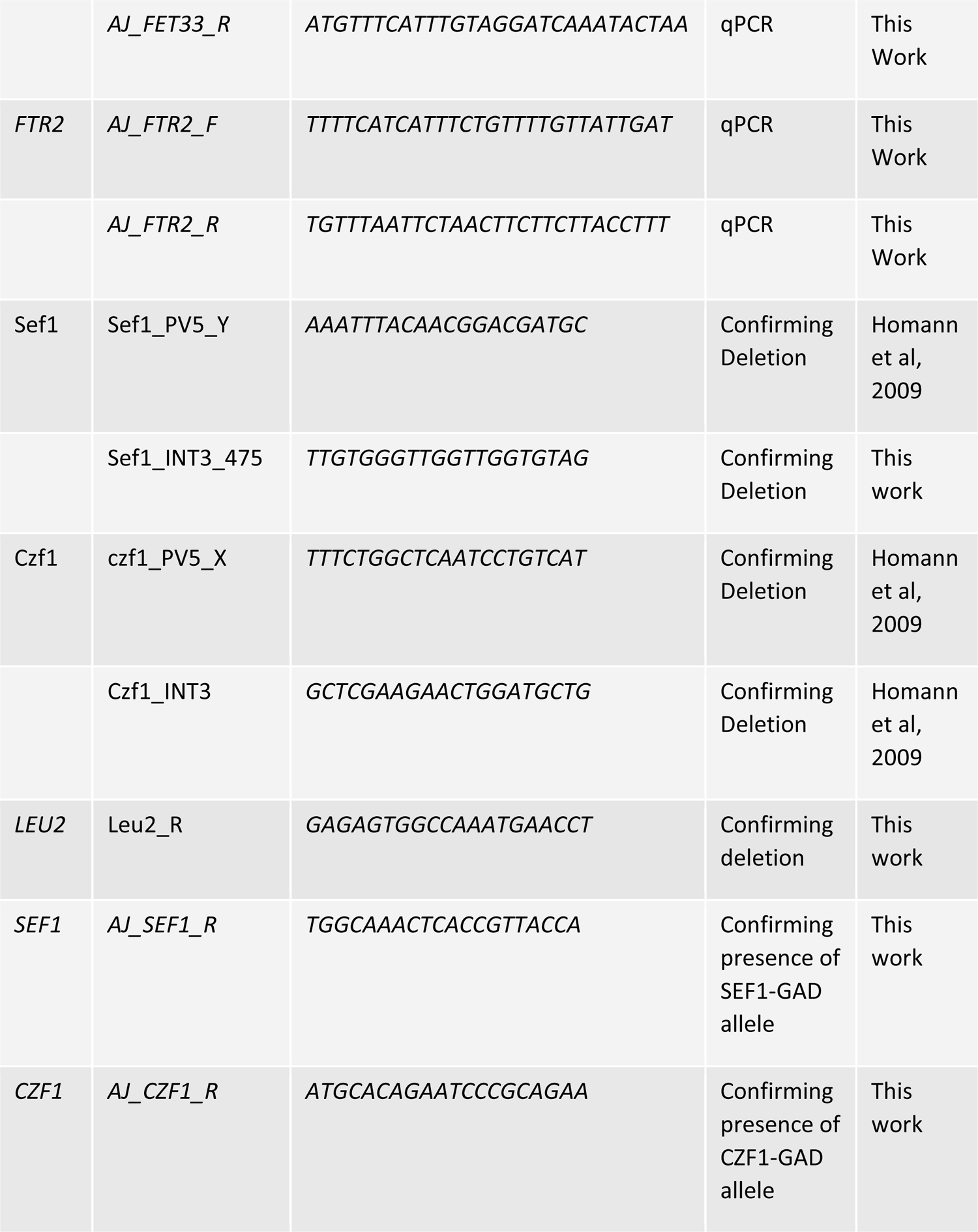
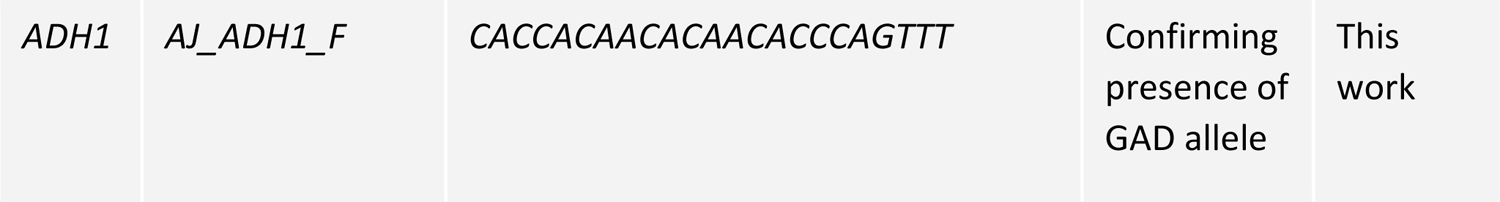
List of Primers Used in This Study

## Acknowledgements

We thank Dr. Joachim Morschhäuser, University of Würzburg for generously providing us with the *CZF1-GAD* and *SEF1-GAD* alleles. Thanks to Albert Tai, Tufts University Genomics Core for help and guidance with the RNAseq processing and data analysis. We also thank Jamison Carey for technical assistance. This research was supported by grant R01AI118898 from the National Institutes of Health (to C.A.K.).

---

**Supplemental Figure S1:**
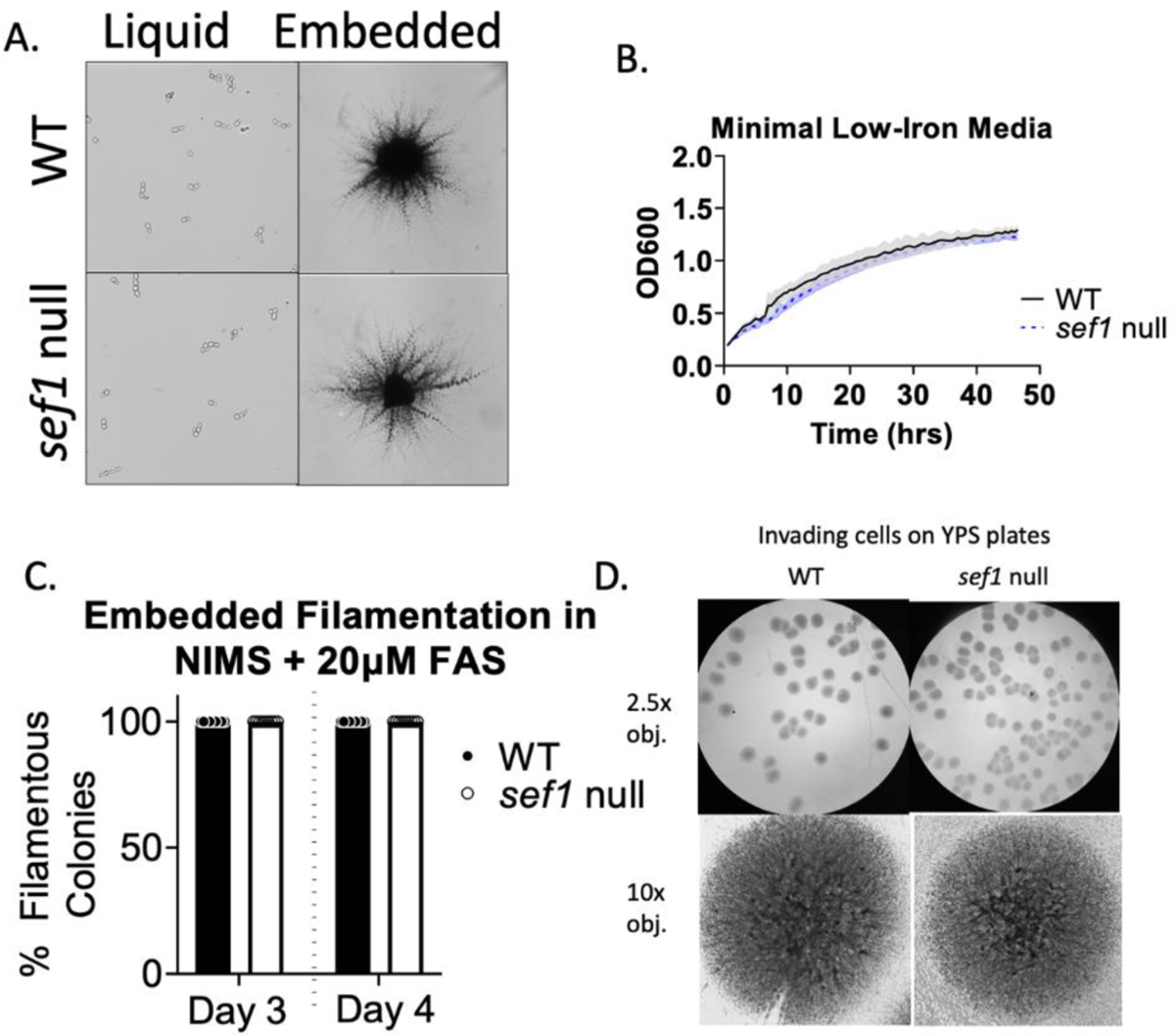
Sef1 is Not Required for Contact-Dependent Filamentation in Low-Iron Media WT and *sef1* null cells were grown overnight in YPD medium at 30°C, then either back diluted to an OD of 0.1 in NIMS media supplemented with 20uM FAS and grown over-day at 30°C or embedded in the same media and allowed to grow for 4 days at 25°C. (A) Brightfield images (4x objective) of WT and *sef1* null cells grown in liquid NIMS (minimal low-iron) media supplemented with 20uM FAS (left) and embedded in NIMS (minimal low-iron) media supplemented with 20uM FAS (right). (B) Growth curve of WT and *sef1* null cells in NIMS + 20uM FAS media at 25°C. Black solid line, WT; gray shading, SD. Blue dotted line, *sef1* null; blue shading, SD. (C) Filamentation of WT and *sef1* null cells grown embedded in NIMS + 20uM FAS media on days 3 and 4 post-embedding. Each point represents 1 biological replicate. 3 experiments with 3 biological replicates per experiment are shown. (D) Images of invading cells left on YPS plates after washing away cells from the agar surface after 4 days of growth. Images were taken using 2.5x and 10x objectives.

**Supplemental Figure S2:**
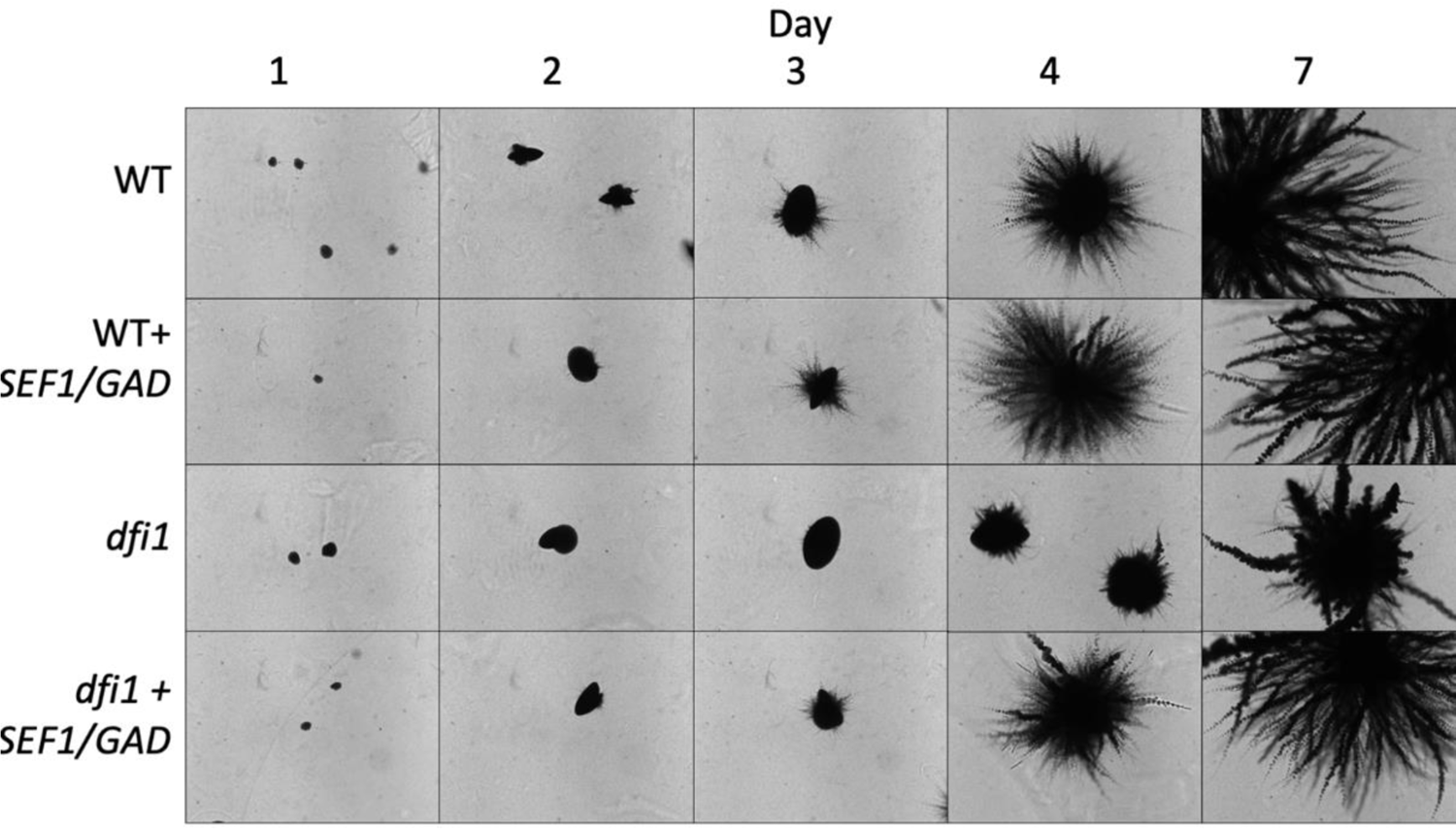
Constitutive Activation of Sef1 in *dfi1* Null mutant WT, *dfi1* null, WT+*SEF1/GAD*, and *dfi1+SEF1/GAD* were embedded in YPS agar media and allowed to grow for 7 days at 25°C. Brightfield images (4x objective) were taken at various times.

**Supplemental Figure S3:**
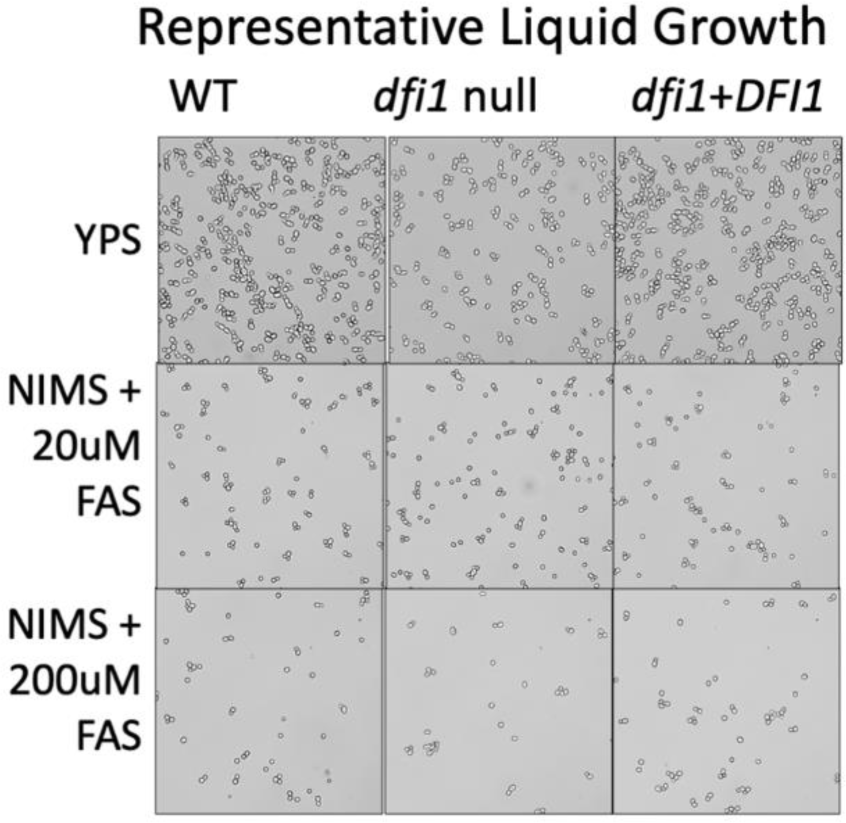
Liquid Low-Iron Media Does Not Induce Filamentation WT, *dfi1* null, or *dfi1+DFI1* cells were grown overnight in YPD medium at 30°C, then back diluted to an OD of 0.1 in liquid YPS or NIMS media supplemented with 20uM or 200uM FAS and grown overnight at 30°C. Brightfield images (4x objective) of WT, *dfi1* null, and *dfi1*+*DFI1* cells grown in liquid rich high-iron or minimal low-iron media.

